## Supplementary Figures for "Sensory Schwann cells set perceptual thresholds for touch and selectively regulate mechanical nociception"

**Extended Data Figures 1-8**

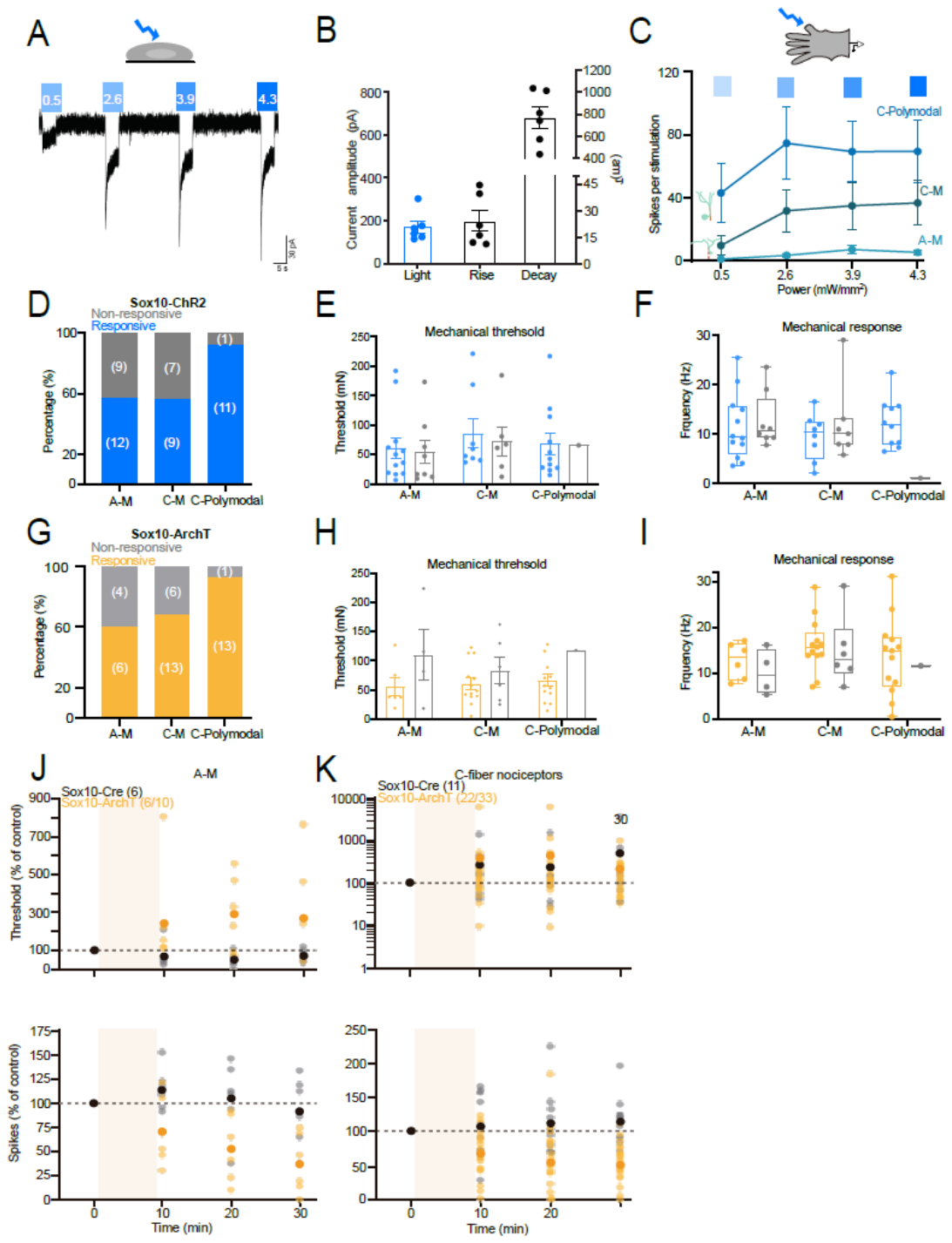

**Extended Data Fig. 1. Light responsive and non-responsive nociceptors.** (**A**) Whole cell patch clamp recordings from sensory Schwann cells in culture isolated from Sox10-Chr2 mice. Inward currents in response to blue light pulses of increasing intensity are shown. (**B**) Current amplitudes, rise times and decay time constants are shown. (**C**) Blue light stimulation of the receptive fields of nociceptors evoked spikes. Note that light intensities of 2.6 mW/mm^2^ and higher evoked maximal responses in the three types of nociceptors measured from Sox10-ChR2 mice (**D**) Proportions of nociceptors of each type that showed activation by blue light in Sox10-ChR2 mice. (**E,F**) The mechanical thresholds and spiking response to a suprathreshold mechanical stimulus (250mN) did not differ between blue light responsive and unresponsive nociceptors. (**G**) Proportions of nociceptors of each type that showed inhibition by yellow light in Sox10-ArchT mice. (**H,I**) The mechanical thresholds and spiking response to a suprathreshold mechanical stimulus (250mN) did not differ between yellow light inhibited and non-inhibited nociceptors. (**J,K)** Changes in threshold and mechanically evoked spiking of individual A-M nociceptors and all C-fiber nociceptors after yellow light exposure in control Sox10-Cre mice (mean black symbol, grey individual neurons) and experimental Sox10-ArchT mice (mean orange circle, yellow individual neurons).

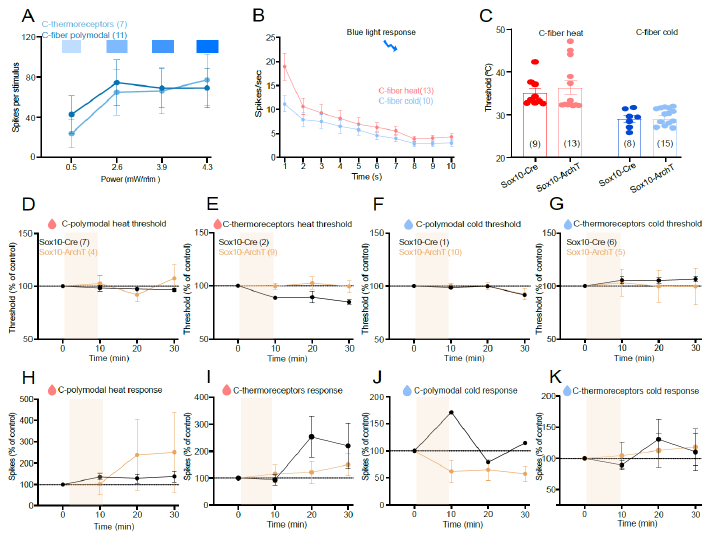

**Extended Data Fig. 2. Schwann cells and thermosensitive C-fibers.** (**A**) Blue light intensities greater than 2.6mW/mm^2^ evoked maximal responses in C-thermoreceptors and polymodal C-fibers. (**B**) Blue light evoked activity was similar in cold and heat sensitive C-fibers (**C**) Thermal thresholds for cold and heat sensitive C-fibers did not differe between Sox10-Cre and Sox10-ArchT mice. (**D-K**) Thermosensitive C-fibers could be classified into four types: heat sensitive polymodal C-fibers and heat sensitive thermoreceptors (**D,E**) and cold sensitive polymodal C-fibers and cold sensitive thermoreceptors (**F,G**). After yellow light exposure neither the heat threshold (top row) nor the spiking response (bottom row) of any of these receptor types was altered in Sox10-ArchT mice compared to Sox10-Cre controls.

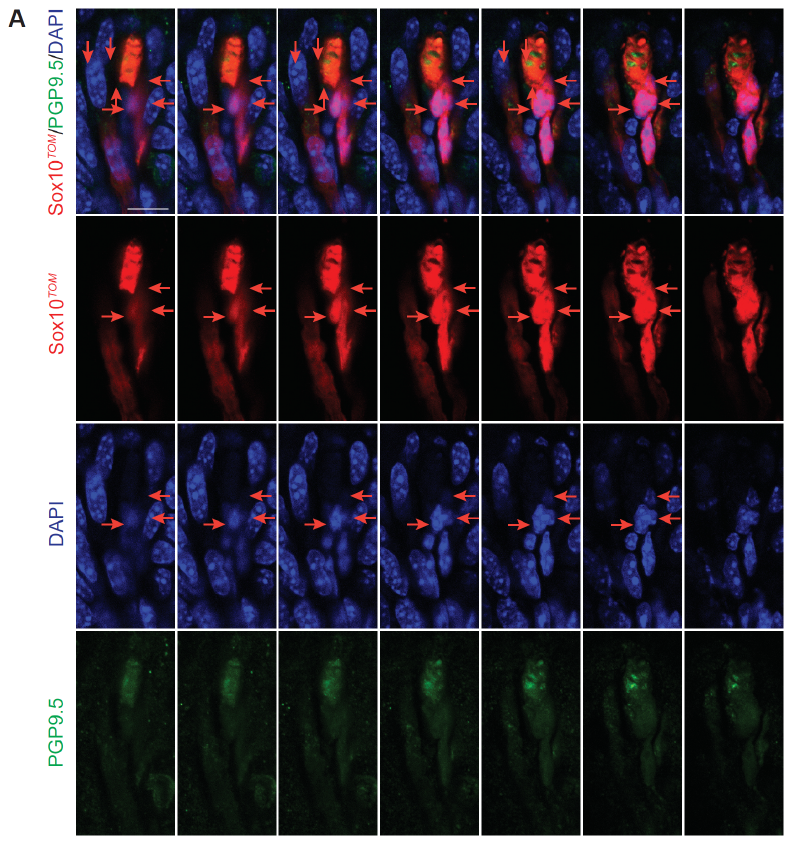

**Extended Data Fig. 3. Targeted/specific recombination in Sox10-Tom mice selectively labels glial cells of Meissner´s corpuscles.** Recombination in glial cells of Meissner corpuscles in Sox10-Tom mice with Tomato (recapitulating *Sox10* expression) and PGP9.5 to label the corpuscle sensory ending and DAPI with Z- stacks (1 μm) images showing the entire corpuscle. Individual channels are used to clearly identify different cells. Arrows point to nuclei from recombined cells in the corpuscle. Scale bar: 20 μm.

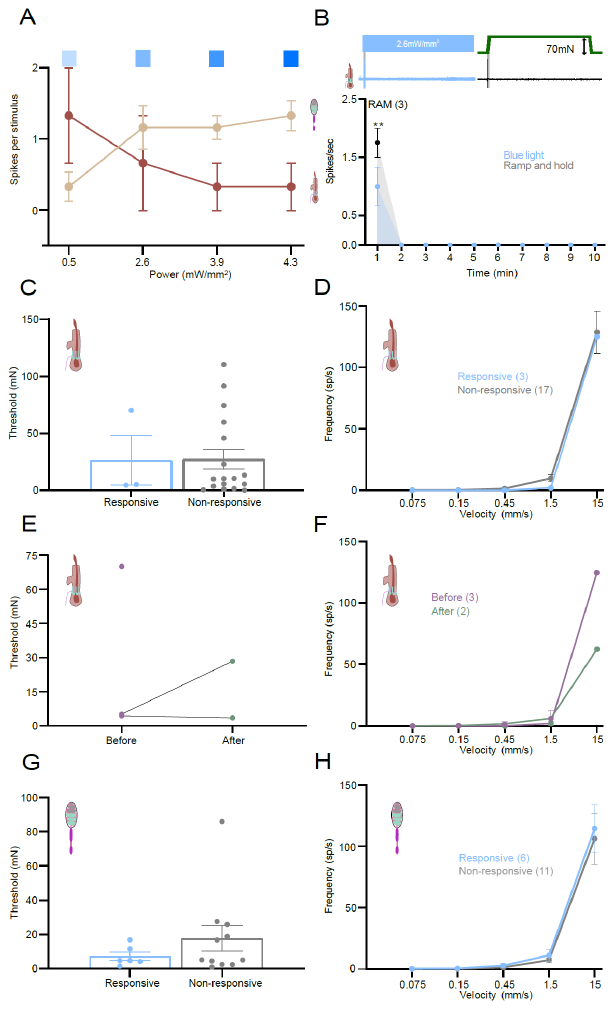

**Extended Data Fig. 4** **Blue light responsive mechanoreceptors in hairy and glabrous skin. (A)** Relationship between light intensity and mechanoreceptor activation **(B)** Action potential firing of RAMs in hairy skin to blue light (blue) and mechanical stimuli (green) in Sox10 ChR2 mice. (**B**) Firing activity in response to mechanical stimuli with varying ramp-velocity of blue light responsive and non-responsive RAMs from Sox10-ChR2 showing similar sensitivities to movement. (**C-D**) Spiking from RAMs exposed to blue light compared to a mechanical stimulus recorded from hairy skin of Sox10-ChR2 mice. (**E-F** RAMs spiking activity before and after blue-light stimulation showing no particular differences in threshold or ramp velocity responses. (G-H)Spiking from RAMs exposed to blue light compared to a mechanical stimulus recorded from glabrous skin of Sox10-ChR2 mice

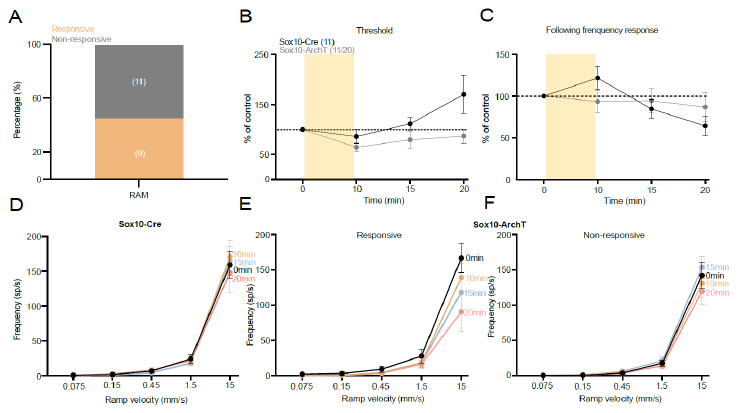

**Extended Data Fig. 5 Physiological properties of non-yellow light responsive RAMs. (A)** Percentage of RAMs responsive (yellow) and non-responsive (gray) to photoinhibition of Sox10^+^ Sensory Schwann cells by yellow light exposure. (**B-C**) Force threshold necessary to evoke the first action potential was observed in RAMs unaffected by yellow light exposure recorded from Sox10-ArchT mice, showing no difference compared to RAM recordings from Sox10-Cre control mice. (**C**) The following frequency from non-affected RAMs before and after yellow light photoinhibition in Sox10-ArchT mice. (**D-**) Firing activity in response to mechanical stimuli with increasing ramp-velocity of RAMs from Sox10-Cre compared to RAMs that were responsive or non-responsive to photoinhibition of Sox10^+^ Sensory Schwann. Movement detection was decreased in RAMs recorded from responsive neurons from Sox10-ArchT mice although this was not statistically significant.

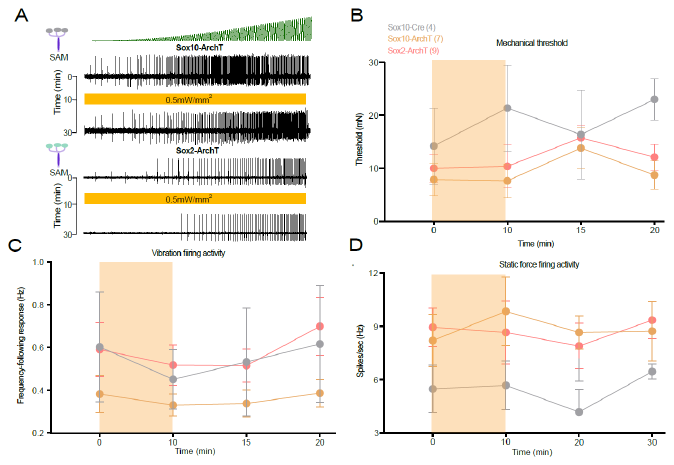

**Extended Data Fig. 6 Slowly adapting mechanoreceptors are not affected by sensory Schwann cell inhibition** (**A**) Examples of spiking of SAMs to a vibration stimulus before and after yellow light exposure in Sox10-ArchT mice (top) and Sox2-ArchT mice (bottom). (**B-D**) Yellow light exposure failed to alter threshold, frequency following or spike frequency of SAMs to mechanical stimuli in Sox10-ArchT and Sox2-ArchT mice. Note that the sample sizes were relatively small so that effects on subsets of SAMs may have been missed.

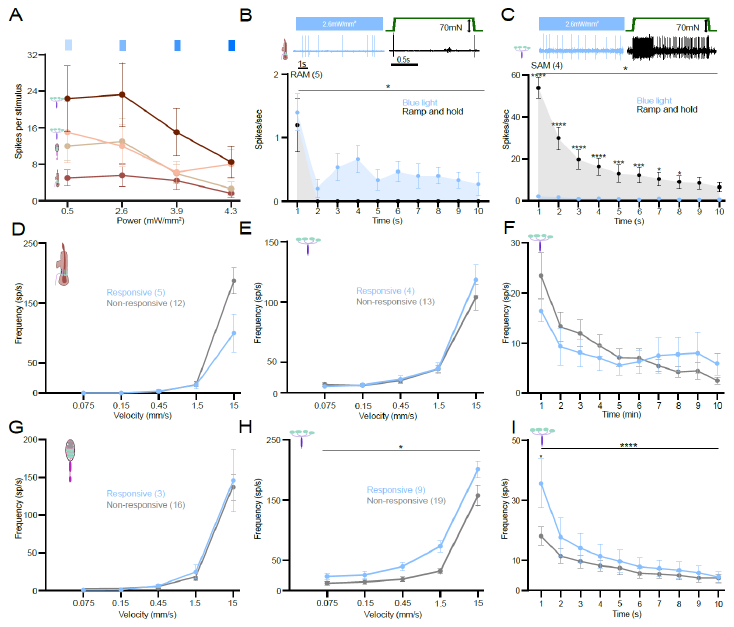

**Extended Data Fig. 7 Activation of RAMs and SAMs by optogenetic stimulation of Sox2 positive Schwann cells. (A)** Low intensity blue light was sufficient to activate SAMs and RAMs in Sox2-ChR2 mice (**B,C**) Blue light stimulation of both hair follicle afferent RAMs and SAMs evoked tonic long latency spiking. Spiking rates in SAMs to blue light were very much lower than responses to mechanical stimulation (**C**). (**D-G**) Mechanosensitivity of blue light responsive and unresponsive RAMs was not different. (**H,I**) Blue light responsive SAMs were significantly more responsive to mechanical stimuli as those that were unresponsive from Sox2-ChR2 mice.

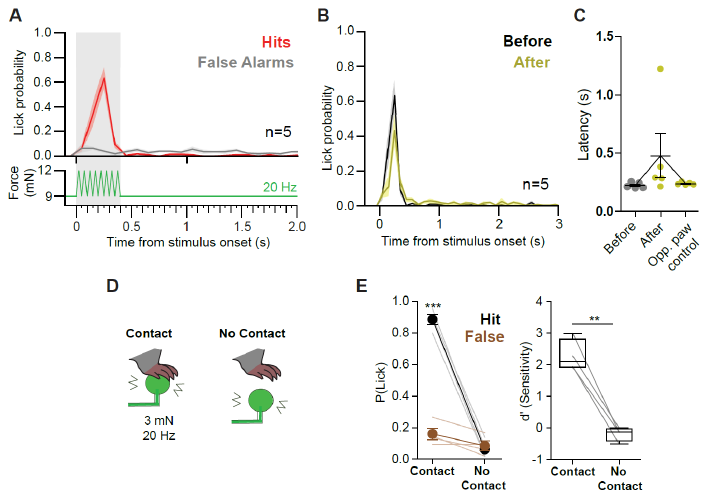

**Extended Data Fig. 8 Mean lick probabilities before and after yellow light exposure** (**A**) (Top) Peri-stimulus time histogram (PSTH) of the lick latencies of Sox10-ArchT mice before optogenetic inhibition (top, n = 5 mice). Licks during stimulus trials are shown in red, and false alarms are in grey. (Bottom) stimulus trial structure. The stimulus consisted on a vibrotactile stimulus of 20 Hz with a baseline force of 9 mN and an amplitude of 3 mN (or 1.5 mN, for some mice). Stimulus duration and window of opportunity (grey box) was was 0.4 s. (**B**) PSTHs of mouse licks during stimulus trials (i.e. hits) before (black) and after the optogenetic inhibition (yellow) (n = 5 mice). (**C**) The median latency to report the stimulus before, after light inhibition and after control light stimulation of the contralateral paw. Optogenetic inhibition did not reach statistical significance for latency change (before vs after, p = 0.0625, Wilcoxon matched-pairs test). (**D**) Cartoon showing control experiment paradigm to check that mouse behavior was not influence by acoustic noise from the stimulator. Mice were trained on the vibrotactile detection task, then the stimulator was moved away from the paw by a few centimeters. (**E**) Hit and false alarm rates were statistically significant when the stimulator was in contact with the skin of Sox10-ArchT mice, but not when the stimulator made no contact (left). Sensitivity (d’) was high (> 1.5) when reporting the stimuli when the stimulator was in contact with the skin, but fell to chance levels (d’ ~ 0) when not in contact with the skin (n = 4 mice).

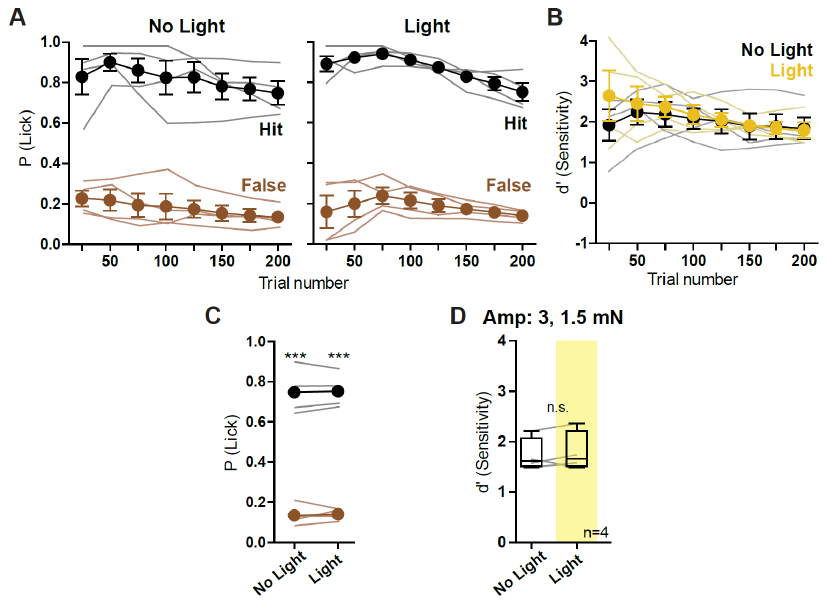

**Extended Data Fig 9. Yellow light stimulation of the forepaw did not impair the detection of vibrotactile stimuli in WT mice. (A)** Percentages of correctly reported trials (hits) and spontaneous licking (false alarms) remained different in the no light as well as the light stimulus behavioral sessions. **(B)** Task performance was not different when comparing the no-light with the light sessions in WT mice. **(C)** Hits were significantly higher than false alarms in both conditions (P < 0.001, Two-way ANOVA with Bonferroni post-hoc, n = 4) **(D)** Performance (sensitivity, d’) was not different between the no-light and light behavioral sessions in WT mice (Wilcoxon matched pairs test, n = 4). The vibrotactile stimulus and the light stimulus parameters were the same as used for Sox10-ArchT mice (shown in Fig. 4 and Extended Data Fig. 6).
